# Consistent decrease in conifer embolism resistance from the stem apex to base resulting from axial trends in tracheid and pit traits

**DOI:** 10.1101/2023.07.21.549999

**Authors:** Zambonini Dario, Savi Tadeja, Rosner Sabine, Petit Giai

## Abstract

Drought-induced embolism formation in conifers is associated with several tracheid and pit traits, which vary in parallel from stem apex to base. We tested whether this axial anatomical variability is associated with a progressive variation in embolism vulnerability along the stem from apex to base.

We assessed the xylem pressure at 50% loss of conductivity (*P50*), the tracheid hydraulic diameter (*Dh*) and mean pit membrane area (*PMA*) on longitudinal stem segments extracted at different distances from the stem apex (*DFA*) in a *Picea abies* and an *Abies alba* tree. In both trees, *Dh* and *PMA* scaled with *DFA*^0.2^. *P50* varied for more than 3 MPa from the treetop to the stem base, according to a scaling of -*P50* with *DFA^-0.2^*. The largest *Dh*, *PMA* and *P50* variation occurred for *DFA*<1.5 m. *PMA* and *Dh* scaled isometrically (exponent *b*=1).

Pit traits vary proportionally with tracheid lumen diameter. Apex-to-base trends in tracheid and pit traits determine a large *DFA*-dependent *P50* variability. Such a *DFA* effect on *P50* did not receive sufficient attention so far, although analysing the relationships *P50* vs. *DFA* is fundamental for the assessment of embolism vulnerability at the individual level.

**Highlights:**

- Conifer embolism vulnerability depends on pit properties, in agreement with published data.
- Pit dimensions increase with tracheid lumen diameter, in agreement with published data
- Tracheid lumen diameter and pit dimensions increase progressively from the stem apex to base, in agreement with published data.
- Xylem vulnerability to embolism formation (*P50*) varies for > 3 MPa from the stem apex to base, with the largest variation occurring within 1.5 m from the stem apex.
- Axial anatomical patterns should be accounted for when analyzing hydraulic properties at individual, intra- and inter-specific scales.

## Introduction

Increasing drought-related tree mortality is a global ecological crisis that demands urgent action from the scientific community. Trees most commonly die from xylem dysfunction by gas-embolism, causing strong limitations to leaf water supply, transpiration, and photosynthesis (Adams et al., 2017; Choat et al., 2018; Hammond, 2020; Savi et al., 2019). Interdisciplinary efforts are necessary to develop risk assessments that identify species-specific vulnerability to drought, enabling a better prediction of climate change impacts on forest ecosystems and global geochemical cycles.

### The mechanism of drought-induced embolism formation

Water moves from roots to leaves following gradients of sub-atmospheric pressure and flows through adjacent vascular conduits. Cohesion and adhesion forces between water molecules and xylem conduit walls maintain the liquid phase of water under tension (Tyree and Zimmermann, 2002). Xylem sap contains dissolved gas bubbles (Schenk et al., 2016) that can cause embolization of the xylem conduit when exceeding a critical bubble diameter (Cochard, 2006). Bordered pits, which connect adjacent vascular elements (vessels, tracheids, and fibers in angiosperms; only tracheids in gymnosperms), serve as a barrier to prevent the entry of large air bubbles into functional elements. The anatomical properties of these pits play a crucial role in determining the vulnerability of xylem to drought-induced embolism formation (Bouche et al., 2014; Lens et al., 2022).

Bordered pits in angiosperms have simple homogeneous membranes with narrow pores, while pits of most conifers have a larger membrane structure with an outer permeable area (margo) and an inner, impermeable disc (torus) larger than the pit aperture. In conifers, when a gas bubble expands and embolizes a tracheid, the pit membranes get aspirated towards the functional conduit due to the pressure difference between the embolized and the functional tracheid (Tyree and Zimmermann, 2002). The margo flexibility and the ratio of torus to pit aperture diameter (i.e., the torus overlap) contribute to a valve effect that seals the pit aperture of the functional tracheid when neighboring tracheids become embolized (Bouche et al., 2014; Delzon et al., 2010).

The estimation of embolism vulnerability is commonly done by measuring the percent loss of xylem hydraulic conductivity (*PLC*) at a specific xylem tension. The contribution of pits to the overall conduit conductance depends on various factors, including the number of pits, their size, and their membrane permeability. Previous studies have suggested that lumen and end-wall resistances contribute approximately 40% and 60%, respectively, to the total xylem resistance, irrespective of conduit size, in both angiosperms and gymnosperms (Sperry et al., 2006). This finding implies the existence of consistent relationships between lumen diameter and pit anatomical traits (Becker et al., 2003; Lazzarin et al., 2016; Schulte, 2012). However, it is important to note that different combinations of traits can result in the same end-wall resistance: in angiosperms, it is primarily influenced by the type of perforation plate, whereas in conifers, by the permeability and area of the margo, as well as the aperture area.

The resistance against drought-induced embolism formation is primarily determined by key pit traits (Bouche et al., 2014; Delzon et al., 2010), leading to a trade-off between hydraulic efficiency and safety (Pittermann et al., 2006). Although this trade-off should be expected to occur mainly at the individual or intra-specific level, it has often been investigated at the interspecific level. One reason why global analyses have failed to provide robust empirical support for the hydraulic efficiency-safety trade-off is the oversight of species-specific combinations of lumen and pit traits (Gleason et al., 2016). Additionally, these types of meta-analyses involve the association of hydraulic and anatomical data obtained using different methods and sampling approaches, such as sampling at fixed branch age or diameter.

These approaches, which are still most commonly used by plant physiologists, do not consider the patterns of variation in anatomical traits that have recently been clearly demonstrated (see below). In the last decade, significant progress has been made to advancing technical capabilities and improving measurement protocols for assessing xylem vulnerability to drought-induced embolism (Cochard et al., 2013; Johnson et al., 2022). However, the contemporary advances in understanding the variability of different xylem anatomical traits and their covariation at individual, intra-specific, and interspecific levels have not received the deserved attention, even though some studies already provided evidence that the xylem vulnerability to drought-induced embolism formation strongly increases axially along the stem from the treetop to the base (Domec and Gartner, 2001).

### The xylem anatomical traits vary axially from the stem apex to base

Recent advancements in sample preparation and computational tools for image analysis allowed research advancements in xylem anatomy, allowing the processing of large amount of data (Prendin et al., 2017).

It has been widely demonstrated that leaves and roots are hydraulically connected through the xylem transport system, with vascular conduits being narrowest at the distal end of the hydraulic path and progressively widening basally along the leaf venation network (Lechthaler et al., 2020) and further down along the outermost, youngest, and longest xylem layer in the stem (Anfodillo et al., 2013) and roots (Prendin et al., 2018). This pattern of widening is consistently repeated every year of growth (Petit et al., 2023) and also occurs radially from the innermost to the outer xylem layers due to continuous height growth during ontogeny (Carrer et al., 2015). Notably, the axial distance from the distal apex (*DFA*) is the best predictor of conduit diameter, according to a power scaling relationship (*Y=a×X^b^*) with an exponent that is nearly invariant at the individual level (Petit et al., 2023; Prendin et al., 2018) and varies only slightly across species (Anfodillo et al., 2013; Koçillari et al., 2021).

Moreover, studies in conifers have reported significant relationships between tracheid lumen diameter and other anatomical features, such as cell wall thickness (Bouche et al., 2014; Prendin et al., 2018; Sperry et al., 2006), tracheid length and number, chamber aperture, torus areas of pits (Becker et al., 2003; Lazzarin et al., 2016; Sperry et al., 2006), torus overlap, margo flexibility (Delzon et al., 2010; Song et al., 2022).

Despite the importance of these allometric relationships for xylem hydraulic efficiency and embolism resistance, they have seldom been investigated and rarely considered in physiological studies. However, there is evidence to suggest that variations in hydraulic traits can be explained by known patterns of xylem anatomical traits, such as the scaling of conduit diameter with *DFA*. Studies focusing on the intra-specific level have shown that embolism vulnerability is lower in terminal twigs than in the stem bole (J. C. Domec et al., 2009; Domec and Gartner, 2001; Rosner et al., 2019b; Spicer and Gartner, 2001). An analysis conducted on *Pseudotsuga menziesii* (Mirb.) Franco reported that the water potential corresponding to 50 % loss of xylem hydraulic conductivity (*P50*) decreased from -3.3 MPa at the stem base to -4.7 MPa at the level of the fifth internode starting from the stem apex (Domec and Gartner, 2001). Furthermore, *P50* was observed to decrease with sapwood depth (Spicer and Gartner, 2001) or being higher at the base of tall trees compared to short trees (Olson et al., 2018; Spicer and Gartner, 2001). In parallel, xylem specific conductivity (*ks*) was reported to increase basally along the stem from the apex downwards and from inner to outer sapwood (Domec et al., 2012; Spicer and Gartner, 2001).

This study aims to provide empirical evidence for how *DFA* affects the variability of anatomical (conduit diameter and pit area) and hydraulic traits (*P50*) in conifer trees. We discuss the importance of using an allometric approach in studies of xylem hydraulics to account for possible axial patterns in anatomical and hydraulic traits, and thus use the scaling parameters to more properly characterize the anatomical and hydraulic traits at the individual and species level.

## Materials and methods

For this study, two mature conifers were selected: a 30-meter silver fir (*Abies alba* Mill.) from a mixed conifer forest located in Asiago, Italy (45° 54’ 6“ N, 11° 29’ 22” E, 1320 m a.s.l.) and a 25-meter Norway spruce (*Picea abies* Karst.) from a pure spruce forest in Enicklberg, Niederösterreich, Austria (48° 14’ 36“ N, 15° 28’ 40” E, 550 m a.s.l.). Both trees were felled in May 2022. Stem discs (19 in the *A. alba* and 25 in the *P. abies* tree) of ∼ 25 cm in length were immediately extracted from at different heights along the stem, and their distance from the stem apex (*DFA*) determined. In order to increase the number of samples within the first 1.5 m of distance from the apex, additional segments were extracted from side branches of the second or third node (in those cases *DFA* referred to the distance from the branch apex). Samples were first debarked, and then sealed in plastic bags with wet paper and transported to the laboratory. For each stem disc, 1-5 sticks of 3 × 5 ×15 cm containing the outermost sapwood rings were obtained from intact and healthy regions with no reaction wood. Apical segments with a diameter < 1 cm were not further processed. All segments were vacuum-sealed and stored at - 20°C. The segments were soaked in distilled water for 24 hours, then split along the fibers to obtain sticks of approximately 14 × 0.8 × 0.8 cm. These sticks contained the three outermost sapwood rings in most cases. The surfaces of the sticks were smoothed with microtome blades. Finally, the sticks were soaked in filtered distilled water purified with silver ions (Micropure, Katadyn Products, Wallisellen, Switzerland) under low-vacuum at room temperature overnight to rehydrate the tissue and refill previously embolized (i.e., air-filled) tracheids (Rosner et al., 2019a, 2018).

### Hydraulic measurements

Vulnerability curves (*VCs*) were assessed by using the air injection technique (Rosner et al., 2021). This technique is the standard method for measuring hydraulic vulnerability of incised sapwood samples from conifer trunks (J.-C. Domec et al., 2009; Domec and Gartner, 2001). Sticks were re-cut under water at both ends with sharp razor blades, connected to a reservoir (at 0.8 m height) containing distilled water with silver ions. The maximum sample hydraulic conductance (*Kmax*) was measured gravimetrically at 8 kPa by collecting sap at the distal end using pre-weighed vials containing a piece of sponge (five vials, 30 seconds interval). Next, the sticks were inserted into a double-ended pressure sleeve (PMS Instruments, Corvallis, OR, USA) and subjected to a pressure of 0.2-0.5 MPa for one minute. The sticks were thus allowed to equilibrate in water for 20 minutes, and the hydraulic conductivity (*K_i_*) was measured again as described above. This process was repeated at increasing pressures, and the percent loss of conductance (*PLC_i_*) was then calculated as:

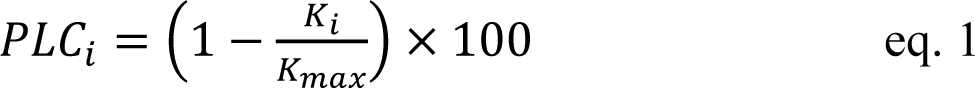

where *K_i_* is the sample conductance measured after the *i*-step of pressurization. The pressure applied in subsequent air injection cycles was gradually increased by steps of 0.5 - 1.0 MPa until *PLC_i_* > 90% was reached.

### Anatomical measurements

Anatomical subsamples of 2 cm were obtained from the center of each stick previously used for hydraulic measurements. These subsamples were cut on cross-section for tracheid lumen diameter measurements and on the longitudinal-radial section for pit measurements using a rotary microtome (LEICA RM2245, Leica Biosystems, Nusslock, Germany) at 10-12 μm, stained with safranine and AstraBlue, and permanently fixed with Eukitt (BiOptica, Milan, Italy). Slides were scanned at 20x with a slide scanner (Axioscan7, Carl Zeiss Microscopy GmbH, Germany). Cross-sections were analyzed with ROXAS (von Arx and Carrer, 2014), which automatically measure the hydraulically weighted mean conduit diameter (*Dh*=*Σd_i_^5^*/*Σd_i_^4^*, where *d_i_* is the diameter of the *i*-conduit, Kolb & Sperry, 1999).

To determine the average surface area of single inter-tracheid pit membranes (*PMA*), radial sections were analyzed using ImageJ software (National Institutes of Health, Bethesda, MD, USA). Specifically, the contour of the pit membrane area was manually drawn on 30-50 pits per sample.

### Statistical analysis and assessment of tracheid vulnerability curve

*VCs* were assessed by fitting *PLC* vs. *P* data by using the fitplc package (Duursma and Choat, 2017) in the R software (R Development Core Team, 2022). For each *VC*, the *P50* value (corresponding to the applied pressure at which *PLC* = 50 %) was extracted.

The allometric relationships between the measured traits were analyzed by using power scaling relationships:

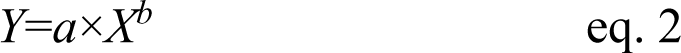

where *a* is the allometric constant and *b* the scaling exponent. Data were first log_10_-transformed to comply with the assumption of normality and homoscedasticity (Zar, 1999), and then fitted with linear regressions, as commonly done in allometric analyses. Eq. 2 was then linearized as:

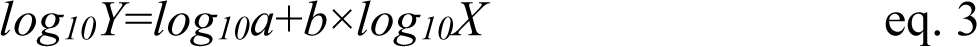

where the allometric constant (*a*) and exponent (*b*) of eq. 2 become the *y*-intercept (=*log_10_a*) and slope (*b*), respectively. The effects of *log_10_DFA*, tree ID and their interaction on hydraulic (*log_10_P50*) and anatomical traits (*log_10_Dh*, *log_10_PMA*) were tested by using linear mixed effects models fitted with restricted maximum likelihood (REML) by using the lme4 R-package (Bates et al., 2015). The best model was chosen based on the Akaike Information Criterion (AIC) (Zuur et al., 2009). Sample ID was used as random factor in all models.

The list of variables used in the hydraulic and anatomical analyses is reported in Table 1.

**Table 1.** List of variables.

| Acronym | Unit | Description |
| --- | --- | --- |
| *DFA* | cm | Axial distance from the stem (or branch) apex |
| *Anatomical variables* |  |  |
| *Dh* | μm | Hydraulically weighted mean tracheid diameter ($Dh = \Sigma d^5 / \Sigma d^4$) |
| *PMA* | μm<sup>2</sup> | Average circular area of tracheid pits |
| *Hydraulic variables* |  |  |
| *PLC* | % | Percent loss of sample conductance |
| *P50* | MPa | Xylem pressure at which PLC=50 % |

## Results

The hydraulically weighted diameter (*Dh*) and the average area of the intertracheid pit membranes (*PMA*) and the water potential inducing the 50 % of conductivity loss (*P50*) showed a large variability down along the stem from apex to base (Fig. 1).

**Figure 1.**
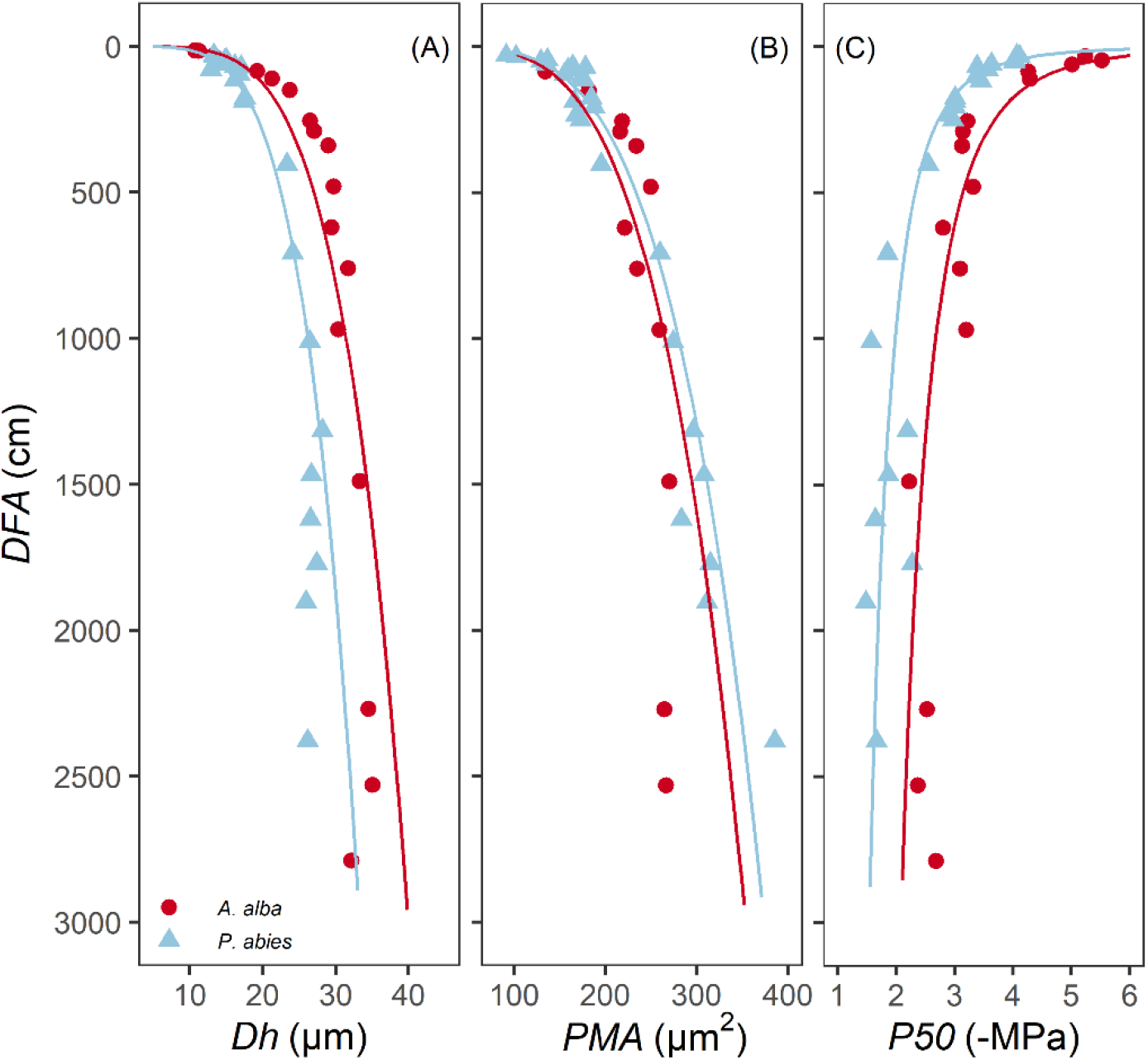
Variation in (A) *Dh*, (B) *PMA* and (C) *P50* with *DFA* in *A. alba* (blue triangles) and *P. abies* (red circles). Fitting lines are according to Table 2.

The anatomical traits *Dh* and *PMA* increased with *DFA* according to power relationships (*Dh*∼*DFA*^0.19^ and *PMA*∼*DFA*^0.23^), whose exponent *b* (eq. 2) was similar in both species (Fig. 2A,B; Table 2A,B). According to our statistical model, the effects of *DFA* and tree *ID* accounted for 92 % of the total *Dh* variance and 88 % of the total *PMA* variance.

**Figure 2.**
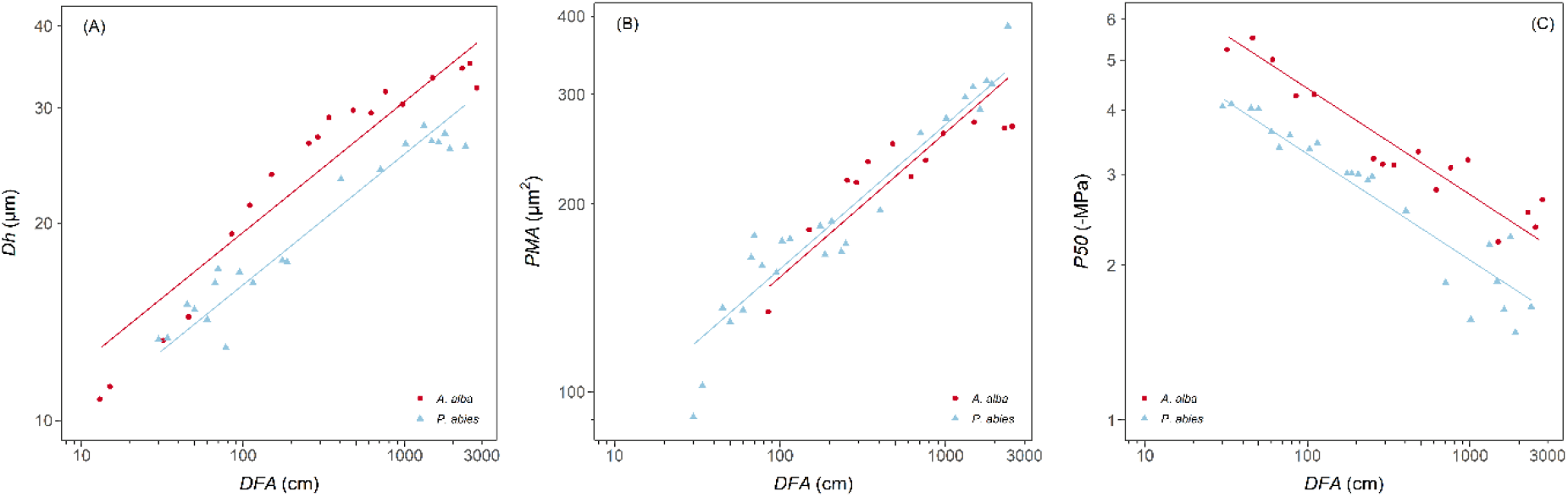
Variation in (A) *Dh*, (B) *PMA* and (C) *P50* with *DFA* in *A. alba* (blue triangles) and *P. abies* (red circles). Fitting lines are according to Table 2. *x* and *y* axis are shown with log_10_ scale.

**Table 2.** Results of the linear mixed-effects models with best AIC predicting the effects of *log_10_DFA* and *Tree ID* on (A) *log_10_Dh*, (B) *log_10_PMA* and (C) *log_10_P50*. *Sample ID* was used as random factor.

|  |  |  |  |  |  |  |
| --- | --- | --- | --- | --- | --- | --- |
| **A** | **Model: $\log_{10}Dh \sim \log_{10}DFA + Tree ID + (Sample ID)$** |  |  |  |  |  |
|  | Fixed effects | Value | Std. error | DF | t value | p value |
|  | y-intercept | 0.89 | 0.03 | 36 | 33.36 | <0.0001 |
| | $\log_{10}DFA$ | 0.20 | 0.01 | 36 | 19.75 | <0.0001 |
|  | *Tree ID (P. abies)* | -0.08 | 0.01 | 36 | -5.74 | <0.0001 |
| | $R^2_{marginal} = 0.92$ | | | | | |
| | $R^2_{conditional} = 0.99$ | | | | | |
| **B** | **Model: $\log_{10}PMA \sim \log_{10}DFA + Tree ID + (Sample ID)$** |  |  |  |  |  |
|  | Fixed effects | Value | Std. error | DF | t value | p value |
|  | y-intercept | 1.72 | 0.04 | 34 | 40.96 | <0.0001 |
| | $\log_{10}DFA$ | 0.23 | 0.01 | 34 | 16.01 | <0.0001 |
|  | *Tree ID (P. abies)* | 0.01 | 0.02 | 34 | 0.71 | 0.4796 |
| | $R^2_{marginal} = 0.88$ | | | | | |
| | $R^2_{conditional} = 0.99$ | | | | | |
| **C** | **Model: $\log_{10}P50 \sim \log_{10}DFA + Tree ID + (Sample ID)$** |  |  |  |  |  |
|  | Fixed effects | Value | Std. error | DF | t value | p value |
|  | y-intercept | 1.06 | 0.03 | 36 | 32.58 | <0.0001 |
| | $\log_{10}DFA$ | -0.21 | 0.01 | 36 | -17.41 | <0.0001 |
|  | *Tree ID (P. abies)* | -0.13 | 0.01 | 36 | -8.51 | <0.0001 |
| | $R^2_{marginal} = 0.90$ | | | | | |
| | $R^2_{conditional} = 0.99$ | | | | | |

*Dh* proportionally increased from apex to base at similar rates in both trees (i.e., same exponent *b*=0.23), but at any *DFA* position in the stem it was significantly narrower in our *P. abies* tree than the *A. alba* tree (i.e., lower *y*-intercept).

Instead, there were no significant differences in the axial pattern of *PMA* along the stem between our sampled trees. *Dh* proportionally increased from apex to base at similar rates in both *P. abies* and *A. alba* trees (i.e., same exponent *b*=0.23), and at any *DFA* position in the stem it was significantly similar between both trees (i.e., similar *y-*intercept).

We found that *PMA* increased proportionally with *Dh* according to a power relationship (*PMA*∼*DFA*^1.2^) (Fig. 3, Table 3), whose exponent is coherent with the exponents characterizing the axial patterns of both *Dh* (*b*=0.19) and *PMA* (*b*=0.23) (note that 0.23/0.19=1.2). Furthermore, *PMA* resulted significantly larger at any observed *Dh* in the *P. abies* than the *A. alba* tree (i.e., higher *y-*intercept: Fig. 3, Table 3).

**Figure 3.**
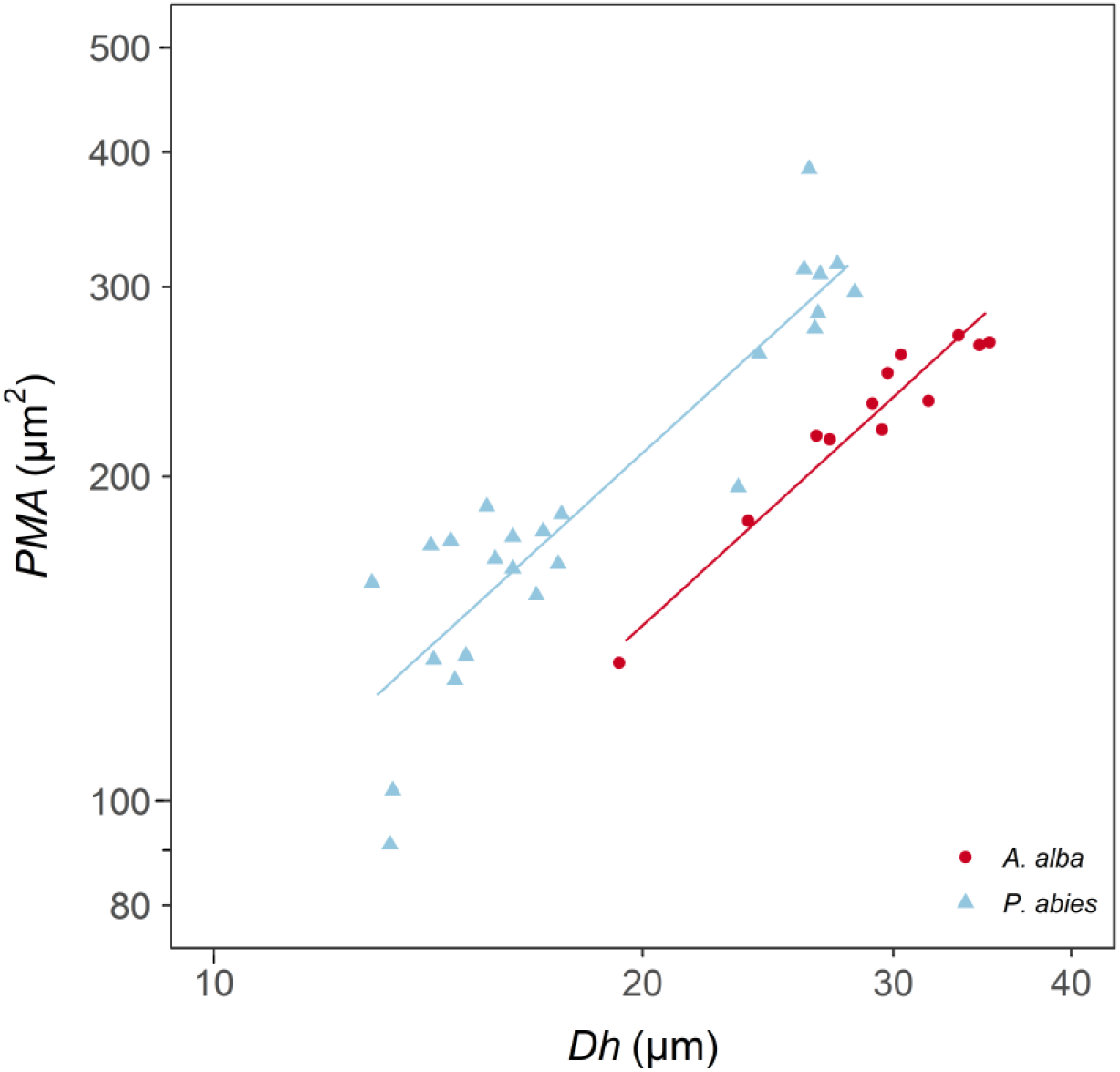
Variation in *PMA* with *Dh* in *A. alba* (blue triangles) and *P. abies* (red circles). Fitting lines are according to Table 3. *x* and *y* axis are shown with log_10_ scale.

**Table 3.** Results of the linear mixed-effects model with best AIC predicting the effects of *log_10_Dh* and *Tree ID* on *log_10_PMA*. *Sample ID* was used as random factor.

| **Model: $\log_{10}PMA \sim \log_{10}Dh + Tree ID + (Sample ID)$** |  |  |  |  |  |
| --- | --- | --- | --- | --- | --- |
| Fixed effects | Value | Std. error | DF | t value | p value |
| y-intercept | 0.60 | 0.13 | 34 | 4.45 | <0.0001 |
| $\log_{10}Dh$ | 1.20 | 0.09 | 34 | 13.25 | <0.0001 |
| *Tree ID (P. abies)* | 0.16 | 0.03 | 34 | 5.98 | <0.0001 |
| $R^2_{marginal} = 0.84$ | | | | | |
| $R^2_{conditional} = 0.98$ | | | | | |

*P50* values ranged from -4.0 to -1.5 MPa for the *P. abies* tree and from -5.5 to -2.2 MPa for the *A. alba* tree. *P50* became progressively less negative along the stem from the apex to base, with ∼1 MPa of the observed variation confined within the most apical 1.5 m of the stem (Fig. 1C and Fig. 2C). This *P50* axial pattern was well described with a power relationship between -*P50* and *DFA* (-*P50*∼*DFA*^-0.21^) (Fig. 2C, Table 2C). -*P50* decreased at similar rates in both sampled trees (i.e., significantly similar exponent, but at any *DFA* position, the *P. abies* tree resulted more vulnerable to air seeding than the *A. alba* tree (i.e., less negative *P50*; higher *y-* intercept). Notably, the effects of *DFA* and tree *ID* explained 90 % of the total *P50* variance. Coherently with the observed relationships between *Dh*, *PMA* and ǀ*P50*ǀ with *DFA*, we found significant power scaling relationships of -*P50* vs. *Dh* and -*P50* vs. *PMA* in both species (Fig. 4, Table 4). *P50* becomes progressively more negative with increasing *Dh* and *PMA* at significantly similar rates in the two sampled trees (-*P50*∼*Dh*^-1.01^, -*P50*∼*PMA*^-0.79^). Tracheids of given *Dh* resulted more vulnerable (i.e., less negative *P50*) in the *P. abies* than the *A. alba* tree (i.e., lower *y-*intercept: Fig. 4A, Table 4A). Seemingly, tracheids of given *PMA* resulted more vulnerable (i.e., less negative *P50*) in the *P. abies* than the *A. alba* tree (i.e., lower *y-* intercept: Fig. 4B, Table 4B).

**Figure 4.**
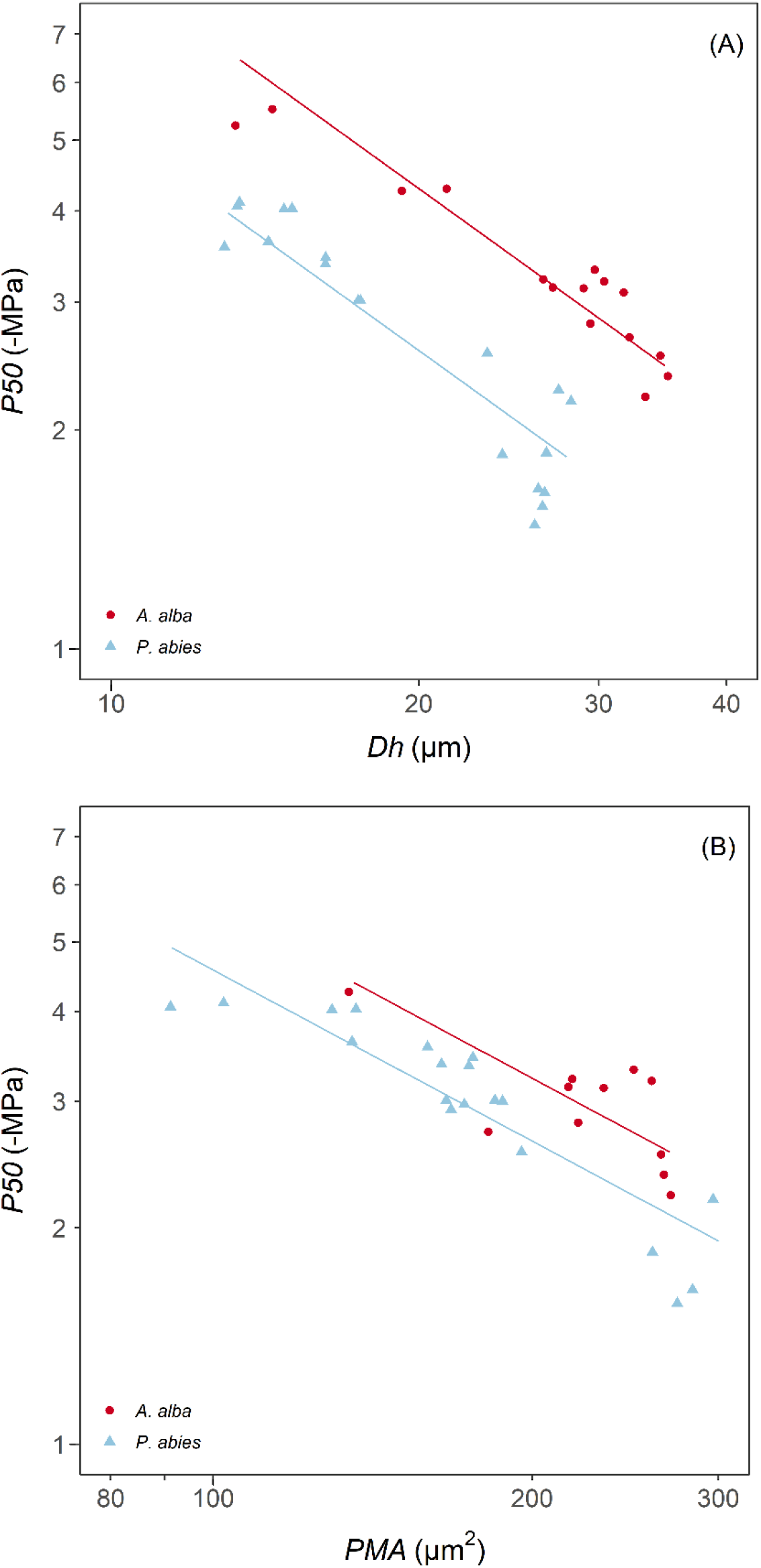
Variation in *P50* with (A) *Dh* and (B) *PMA* in *A. alba* (blue triangles) and *P. abies* (red circles). Fitting lines are according to Table 4. *x* and *y* axis are shown with log_10_ scale.

**Table 4.** Results of the linear mixed-effects models with best AIC predicting the effects of *Tree ID* and (A) *log_10_Dh* and (B) *log_10_PMA* on *log_10_P50*. *Sample ID* was used as random factor.

| **A** | **Model: $\log_{10}P50 \sim \log_{10}Dh + species + (ID)$** |  |  |  |  |  |
| --- | --- | --- | --- | --- | --- | --- |
|  | Fixed effects | Value | Std. error | DF | t value | p value |
|  | y-intercept | 1.95 | 0.11 | 31 | 18.01 | <0.0001 |
| | $\log_{10}Dh$ | -1.01 | 0.08 | 31 | -13.36 | <0.0001 |
|  | *Tree ID (P. abies)* | -0.22 | 0.02 | 31 | -10.31 | <0.0001 |
| | $R^2_{marginal} = 0.86$ | | | | | |
| | $R^2_{conditional} = 0.98$ | | | | | |
| **B** | **Model: $\log_{10}P50 \sim \log_{10}PMA + species + (ID)$** |  |  |  |  |  |
|  | Fixed effects | Value | Std. error | DF | t value | p value |
|  | y-intercept | 2.33 | 0.17 | 31 | 13.72 | <0.0001 |
| | $\log_{10}PMA$ | -0.79 | 0.07 | 31 | -11.02 | <0.0001 |
|  | *Tree ID (P. abies)* | -0.09 | 0.02 | 31 | -3.92 | <0.0001 |
| | $R^2_{marginal} = 0.79$ | | | | | |
| | $R^2_{conditional} = 0.97$ | | | | | |

It can be summarized that (i) the power scaling relationships of *Dh*, *PMA* and *P50* with *DFA* showed no differences in the scaling exponent between the two sampled trees, (ii) at any *DFA* position each trait differed between the sampled trees by a constant factor equal to the ratio of their allometric constant *a* (eq. 2). In practice, at any *DFA* position along the stem, tracheids of the *P. abies* tree had 20 % narrower *Dh*, similar *PMA* and 35 % less negative *P50*.

## Discussion

The study revealed significant axial patterns in all analyzed traits (*Dh*, *PMA* and *P50*). The increase in *Dh* and *PMA* from stem apex to base aligned with previous anatomical analyses on conifers (Anfodillo et al., 2012, 2006; Jyske and Hölttä, 2015; Lazzarin et al., 2016; Lintunen et al., 2010; Petit et al., 2023, 2011). Besides, our hydraulic measurements provided new empirical evidence of a consistent axial pattern in *P50*, becoming less negative with increasing distance from the stem apex (*DFA*) to the base.

### The functional allometry of tracheid and pit traits

All traits exhibited allometric scaling with *DFA*, following power relationships (eq. 2). These relationships are commonly found in all living organisms, representing fundamental functional properties (West et al., 1999, 1997). When one trait varies, several others change proportionally to maintain essential functional properties. The scaling exponent (*b*) describes the relative variation in the *Y* trait for a given change in the *X* trait. For example, trees maintain their upright position by increasing stem diameter (*D*) in precise proportion to the increase in tree height (*H*) to ensure mechanical stability (resulting in the universal scaling of *H*∼*D*^2/3^: McMahon & Kronauer, 1976; Niklas, 1995).

Among the traits analyzed in this study, the hydraulically weighted tracheid lumen diameter (*Dh*) received the most attention. In our trees, *Dh* increased with *DFA*^0.2^ (scaling exponent *b*=0.2), consistent with previous studies on axial *Dh* variation along the youngest / outermost xylem ring (Anfodillo et al., 2012, 2006; Jyske and Hölttä, 2015; Lintunen and Kalliokoski, 2010; Petit et al., 2023, 2011). This axial pattern has been traditionally referred to as “conduit tapering” (the progressive reduction in lumen diameter from the stem base towards the apex: Schulte, 2012; Pfautsch *et al*., 2018). However, the axial *Dh* variation is strongly *DFA*-dependent, since such a pattern was found to be rigidly reiterated at every year of growth (i.e., the same pattern can be found along each annual ring from the ring apex to the stem base: Prendin *et al*., 2018; Petit *et al*., 2023). This pervasive *DFA*-dependent *Dh* pattern, better termed “widening”, has been hypothesized to contribute essential hydraulic properties to the xylem architecture. From one hand, larger conduits towards the stem base would contribute very little to the total hydraulic resistance to water transport (Becker et al., 2000; Petit and Anfodillo, 2009), thus effectively contributing to maintain an efficient leaf water supply while growing taller (Petit et al., 2010; West et al., 1999). On the other, increasingly conductive conduits from the stem apex to base would strongly affect the shape of the water potential (*Ψ*) gradient from the stem apex to base, with the lowest (i.e., most negative) *Ψ* concentrated within a short distance from the apex (Lechthaler et al., 2020).

Notably, the above conditions received indirect empirical support from studies reporting very limited variation in the contribution of pit resistance to the total conduit (either vessels in angiosperms and tracheids in conifers) resistance with increasing conduit diameter (Domec et al., 2006; Petit et al., 2008; Pittermann et al., 2006).

### Tracheid-pit allometry: the inflatable balloon effect

The development of tracheids can be compared to inflatable balloons that expand in volume as the internal pressure (turgor) increases, stretching the cell wall structures (Cosgrove, 1986). Consistent with this hypothesis, tracheid diameter and different pit traits have been reported to be highly correlated both at the individual (Lazzarin et al., 2016; Schulte, 2012) and interspecific level (Delzon et al., 2010; Hacke et al., 2004).

Our measurements of *PMA* and *Dh* at different positions along the stem provided new evidence that tracheid and pit traits tightly vary in tandem along the stem from apex to base (Lazzarin et al., 2016; Schulte, 2012). Both *PMA* and *Dh* varied with *DFA^b^*, with the scaling exponent for both relationships and in both *A. alba* and *P. abies* tree being *b*∼0.2, thus very similar to the same scaling relationships reported for the giant *Sequoiadendron giganteum* (Lindl.) J.Buchh. tree analyzed by Lazzarin *et al*. (2016).

Furthermore, Lazzarin *et al*. (2016) reported highly significant *DFA*-dependent trends not only for *PMA*, but also for torus area (*TA*) and pit aperture area (*PA*). Since these traits are typically highly correlated (Delzon et al., 2010; Hacke et al., 2004), it seems highly likely that also in our analysed trees *TA* and *PA* conformed to these relationships with *PMA*, and therefore progressively increased with *DFA* (additional indirect evidence is discussed below).

The regulation of the tracheid lumen diameter, as well as pit traits, is likely associated to the duration of cell elongation, which progressively increases from the stem apex to base (Anfodillo et al., 2012).

### Increasing embolism vulnerability from the stem apex to base

While it is widely acknowledged that pit traits (*PMA*, *TA*, and *PA*) strongly influence the resistance against embolism formation through air seeding via pit membranes (Bouche et al., 2014; Delzon et al., 2010; Hacke et al., 2004), the growing body of empirical evidence demonstrating strong and consistent (almost universal) anatomical patterns dependent on *DFA* has not adequately motivated the scientific community to investigate the impact of *DFA* on xylem hydraulic properties, such as embolism vulnerability.

We observed a tremendous variation in *P50* (>3 MPa) along the stem of both the *P. abies* and *A. alba* trees. This axial variability is twice as large as that reported for *Pseudotsuga menziesii* (Domec and Gartner, 2001). However, it is important to note that Domec & Gartner did not sample segments between the stem apex and the fifth internode, which could be located at a distance of over 1 meter from the stem apex.

The minimum *P50* values of both trees were in line with those found in the literature and obtained from *VCs* of apical branches (<1 m) measured with different techniques (bench dehydration and hydraulic measurements, acoustic, centrifugation, etc.) (Choat et al., 2012; Feng et al., 2021). Even if sometimes criticised, the air injection method is well established for analyses on conifers and short-vessel species, and several studies showed good agreements between the air injection and other methodical approaches (Rosner et al., 2019a).

The axial pattern of *P50* strongly depended on *DFA*, following a power trajectory similar to *Dh* and *PMA* (ǀ*b*ǀ∼0.2). However, it also resembled the trajectory of *TA* and *PA*, indicating an inflating balloon effect on tracheid growth (discussed above). *P. abies* exhibited less negative *P50* (Fig. 2C, Table 2C), narrower *Dh* (Fig. 2A, Table 2A) and similar *PMA* (Fig. 2B, Table 2B) compared to *A. alba* along the stem. It would be a mistake to interpret this as the *P. abies* tree being more vulnerable due to its narrower tracheids. Instead, this comparison supported the hypothesis that the *Dh-P50* relationship lacks causality (Lens et al., 2022). Tracheids with a given *PMA* showed higher vulnerability to gas-embolism in the *P. abies* than in the *A. alba* tree (lower *y*-intercept: Table 4B, Fig. 4B). However, both trees exhibited similar *PMA* at any *DFA* position (Table 2B, Fig. 2B), while *P50* differed (Table 2C, Fig. 2C). This can be explained by other pit traits, such as torus area (*TA*) and the aperture area (*PA*), which likely varied proportionally with *Dh* and *PMA* in our trees (Lazzarin et al., 2016). Most likely, at any *DFA* and *PMA*, the *P. abies* tree resulted more vulnerable than the *A. alba* tree due to its narrower *TA* and/or larger *PA*, corresponding to a less efficient torus overlap (Hacke and Jansen, 2009). Indeed, the correlation between different anatomical traits (tracheid diameter, pit membrane area, torus area and aperture area) is typically very high (Bouche et al., 2014; Delzon et al., 2010), in agreement with our data and the findings of Lazzarin *et al*. (2016) on axial trends in anatomical traits.

The within-tree *P50* variability reported in this study for only two trees of similar species corresponded to approximately 60% of the *P50* variability among many gymnosperm species reported in a previous study (Choat et al., 2012).

To ensure comparability of xylem hydraulic analyses, it is crucial to consider the potential *DFA* effects on lumen diameter, pit traits, and hydraulic characteristics.

Xylem hydraulic data, such as *P50* and xylem specific conductivity (*ks*), are commonly obtained by sampling stem / branch segments with standardized age (i.e., number of rings) or diameter, and specific length (depending on methods), thus they do not account for *DFA* effects. Therefore, if *DFA* is unknown, accurate comparability of published data is quite challenging.

### Towards the allometric scaling of plant hydraulics

The observed increase in *P50*, becoming less negative, along the stem from apex to base is in line with the predicted pattern based on the axial variation in tracheid and pit traits mentioned earlier.

Removing the DFA effect from hydraulic measurements is not a straightforward task. In our trees -*P50* scaled with *DFA*^-0.2^. However, our limited data cannot exclude that *b* may be higher or lower than -0.2 depending on species and/or environmental conditions. Therefore, sampling at fixed *DFA* does not entirely eliminate the *DFA* effects on *P50*, unless the *b*=-0.2 will be demonstrated to be universal in plants.

Instead, considering the axial variability in anatomical (e.g., conduit diameter and pit traits) and hydraulic (e.g., xylem specific conductivity, *P50*) traits would help account for the *DFA* effects. Applying our allometric approach, we could determine that the analyzed *P. abies* tree was 35 % more vulnerable than the *A. alba* tree: since -*P50*∼*DFA-0.2* (similar exponent *b*=0.2), at any *DFA* position along the stem the ratio of *P50* between the two trees was equal to the ratio of their allometric constant *a* (eq.1) (*a_P.abies_/a_A.alba_*=1.35). This relationship can be visually represented as parallel lines on a log-log graph (using either log-transformed data or a linear scale with logarithmic axes) (cf. case A vs. case B in Fig. 5: same slope and different *y*-intercept).

**Figure 5.**
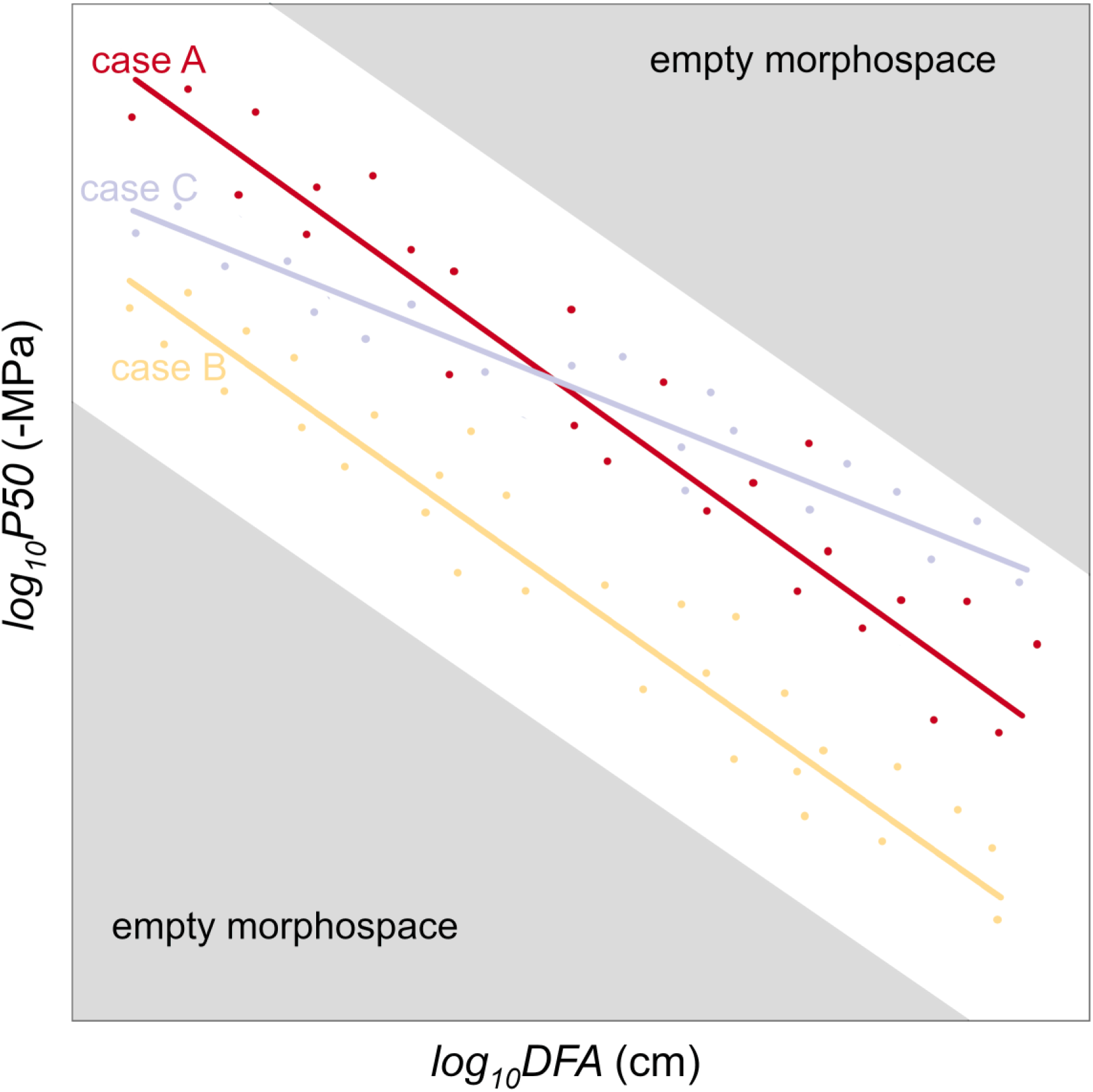
How allometric trajectories can be used for studying the variability of *P50* (or other hydraulic traits) in different contexts. The base case A represents the *P50* variation with *DFA* at the single individual level. Compared to case A, the lower *y-*intercept of case B indicate a lower resistance against drought-induced embolism formation; the lower *y-*intercept and higher slope of case C indicate a lower embolism resistance in the upper stem and a higher resistance in the lower stem. Convergence towards a similar scaling of *P50* with *DFA* could be the result of selection eliminating variants with too high *P50* towards the apex (where the xylem tensions are the highest due to the proximity to the transpiring leaves), and too low tracheid conductivity towards the base (determining hydraulic limitations to leaf water supply).

In cases where two individuals would have different values for *a* and *b* (eq. 2), resulting in distinct *y*-intercepts and slopes on log-log representation (eq. 2) (cases A and C of Fig. 5), measuring *P50* at a single position in the branch would lead to one individual (case A) appearing less vulnerable if samples were taken towards the apex, or more vulnerable if samples were taken towards the base.

Further investigations are needed to test the consistency and generality of the axial scaling of embolism vulnerability (*P50*) with *DFA*. This approach could provide a new perspective on studying plant functional traits, including anatomical and hydraulic characteristics. Empirical evidence increasingly supports the notion of species converging towards a universal scaling of conduit diameter (*Dh*) with *DFA* (Anfodillo et al., 2013; Kiorapostolou and Petit, 2019; Lechthaler et al., 2019; Olson et al., 2018; Petit et al., 2023). Additionally, there is growing evidence of similar scaling relationships between pit traits and *DFA* (this study, Schulte, 2012; Lazzarin *et al*., 2016). It is reasonable to assume that evolution has favored variants that strike a balance between safety and efficiency, eliminating those with vulnerable conduits towards the apex (where xylem pressures are most negative due to proximity to transpiring leaves) and those with low conductive vascular elements towards the base (which would limit leaf water supply and gas exchange efficiency). This can be visualized as empty morphospaces in Fig. 8.

## Conclusion

This study presents novel empirical evidence that supports the concept of xylem tracheids behaving like inflatable balloons, with all xylem anatomical traits strongly influenced by their distance from the apex. This results in significant variation in *P50* along the stem, from the apex to the base.

We strongly emphasize the importance for researchers investigating xylem hydraulics to carefully consider the potential impact of axial distance from the apex on traits such as embolism vulnerability. Accounting for this effect is crucial for obtaining accurate measurements, enabling robust analysis and comparison of individuals, populations, and species. Ultimately, this knowledge will enhance our ability to predict the impacts of climate change on plant communities.

## Statement of author contribution

GP SR and TS designed the research. DZ and TS performed the measurements. DZ performed the statistical analyses. DZ and GP wrote the first draft of the manuscript. All authors contributed critically to the final draft of the manuscript.

## Acknowledgements

We warmly thank Khristina Zagudaeva, Christian Rebeschini, Petr Zabransky and Daniel Gröticke for technical assistance. DZ and GP were supported by Fondazione Cariparo and University of Padua (DOR2202355/22), respectively.

## Bibliography

1. Adams, H.D., Zeppel, M.J.B., Anderegg, W.R.L., Hartmann, H., Landhäusser, S.M., Tissue, D.T., Huxman, T.E., Hudson, P.J., Franz, T.E., Allen, C.D., Anderegg, L.D.L., Barron-Gafford, G.A., Beerling, D.J., Breshears, D.D., Brodribb, T.J., Bugmann, H., Cobb, R.C., Collins, A.D., Dickman, L.T., Duan, H., Ewers, B.E., Galiano, L., Galvez, D.A., Garcia-Forner, N., Gaylord, M.L., Germino, M.J., Gessler, A., Hacke, U.G., Hakamada, R., Hector, A., Jenkins, M.W., Kane, J.M., Kolb, T.E., Law, D.J., Lewis, J.D., Limousin, J.-M., Love, D.M., Macalady, A.K., Martínez-Vilalta, J., Mencuccini, M., Mitchell, P.J., Muss, J.D., O’Brien, M.J., O’Grady, A.P., Pangle, R.E., Pinkard, E.A., Piper, F.I., Plaut, J.A., Pockman, W.T., Quirk, J., Reinhardt, K., Ripullone, F., Ryan, M.G., Sala, A., Sevanto, S., Sperry, J.S., Vargas, R., Vennetier, M., Way, D.A., Xu, C., Yepez, E.A., McDowell, N.G., 2017. A multi-species synthesis of physiological mechanisms in drought-induced tree mortality. Nat. Ecol. Evol. 1, 1285–1291. https://doi.org/10.1038/s41559-017-0248-x

2. Anfodillo, T., Carraro, V., Carrer, M., Fior, C., Rossi, S., 2006. Convergent tapering of xylem conduits in different woody species. New Phytol. 169, 279–290. https://doi.org/10.1111/j.1469-8137.2005.01587.x

3. Anfodillo, T., Deslauriers, A., Menardi, R., Tedoldi, L., Petit, G., Rossi, S., 2012. Widening of xylem conduits in a conifer tree depends on the longer time of cell expansion downwards along the stem. J. Exp. Bot. 63, 837–845. https://doi.org/10.1093/jxb/err309

4. Anfodillo, T., Petit, G., Crivellaro, A., 2013. Axial conduit widening in woody species: A still neglected anatomical pattern. IAWA J. 34, 352–364. https://doi.org/10.1163/22941932-00000030

5. Bates, D., Mächler, M., Bolker, B., Walker, S., 2015. Fitting Linear Mixed-Effects Models Using lme4. J. Stat. Softw. 67, arXiv:1406.5823. https://doi.org/10.18637/jss.v067.i01

6. Becker, P., Gribben, R.J., Lim, C.M., 2000. Tapered conduits can buffer hydraulic conductance from path-length effects. Tree Physiol. 20, 965–967. https://doi.org/10.1093/treephys/20.14.965

7. Becker, P., Gribben, R.J., Schulte, P.J., 2003. Incorporation of transfer resistance between tracheary elements into hydraulic resistance models for tapered conduits. Tree Physiol. 23, 1009–1019.

8. Bouche, P.S., Larter, M., Domec, J.-C., Burlett, R., Gasson, P., Jansen, S., Delzon, S., 2014. A broad survey of hydraulic and mechanical safety in the xylem of conifers. J. Exp. Bot. 65, 4419–4431. https://doi.org/10.1093/jxb/eru218

9. Carrer, M., Von Arx, G., Castagneri, D., Petit, G., 2015. Distilling allometric and environmental information from time series of conduit size: The standardization issue and its relationship to tree hydraulic architecture. Tree Physiol. https://doi.org/10.1093/treephys/tpu108

10. Choat, B., Brodribb, T.J., Brodersen, C.R., Duursma, R.A., López, R., Medlyn, B.E., 2018. Triggers of tree mortality under drought. Nature 558, 531–539. https://doi.org/10.1038/s41586-018-0240-x

11. Choat, B., Jansen, S., Brodribb, T.J., Cochard, H., Delzon, S., Bhaskar, R., Bucci, S.J., Feild, T.S., Gleason, S.M., Hacke, U.G., Jacobsen, A.L., Lens, F., Maherali, H., Martínez-Vilalta, J., Mayr, S., Mencuccini, M., Mitchell, P.J., Nardini, A., Pittermann, J., Pratt, R.B., Sperry, J.S., Westboy, M., Wright, I.J., Zanne, A.E., 2012. Global convergence in the vulnerability of forests to drought. Nature 491, 752–755. https://doi.org/10.1038/nature11688

12. Cochard, H., 2006. Cavitation in trees. Comptes Rendus Phys. 7, 1018–1026. https://doi.org/10.1016/j.crhy.2006.10.012

13. Cochard, H., Badel, E., Herbette, S., Delzon, S., Choat, B., Jansen, S., 2013. Methods for measuring plant vulnerability to cavitation: a critical review. J. Exp. Bot. 64, 4779–4791. https://doi.org/10.1093/jxb/ert193

14. Cosgrove, D., 1986. Biophysical control of plant cell growth. Annu. Rev. Plant Physiol. 37, 377–405. https://doi.org/10.1146/annurev.pp.37.060186.002113

15. Delzon, S., Douthe, C., Sala, A., Cochard, H., 2010. Mechanism of water-stress induced cavitation in conifers: bordered pit structure and function support the hypothesis of seal capillary-seeding. Plant. Cell Environ. 33, 2101–2111. https://doi.org/10.1111/j.1365-3040.2010.02208.x

16. Domec, J.-C., Lachenbruch, B., Pruyn, M.L., Spicer, R., 2012. Effects of age-related increases in sapwood area, leaf area, and xylem conductivity on height-related hydraulic costs in two contrasting coniferous species. Ann. For. Sci. 69, 17–27. https://doi.org/10.1007/s13595-011-0154-3

17. Domec, J.-C., Noormets, A., King, J.S., Sun, G.E., McNulty, S.G., Gavazzi, M.J., Boggs, J.L., Treasure, E.A., 2009. Decoupling the influence of leaf and root hydraulic conductances on stomatal conductance and its sensitivity to vapour pressure deficit as soil dries in a drained loblolly pine plantation. Plant. Cell Environ. 32, 980–991. https://doi.org/10.1111/j.1365-3040.2009.01981.x

18. Domec, J.C., Gartner, B.L., 2001. Cavitation and water storage capacity in bole xylem segments of mature and young Douglas-fir trees. Trees - Struct. Funct. 15, 204–214. https://doi.org/10.1007/s004680100095

19. Domec, J.C., Lachenbruch, B., Meinzer, F.C., 2006. Bordered pit structure and function determine spatial patterns of air-seeding thresholds in xylem of Douglas-fir (Pseudotsuga menziesii; Pinaceae) trees. Am. J. Bot. 93, 1588–1600.

20. Domec, J.C., Warren, J.M., Meinzer, F.C., Lachenbruch, B., 2009. Safety factors for xylem failure by implosion and air-seeding within roots, trunks and branches of young and old conifer trees. IAWA J. 30, 101–120.

21. Duursma, R., Choat, B., 2017. fitplc - an R package to fit hydraulic vulnerability curves. J. Plant Hydraul. 4, e002. https://doi.org/10.20870/jph.2017.e002

22. Feng, F., Losso, A., Tyree, M., Zhang, S., Mayr, S., 2021. Cavitation fatigue in conifers: a study on eight European species. Plant Physiol. 186, 1580–1590. https://doi.org/10.1093/plphys/kiab170

23. Gleason, S.M., Westoby, M., Jansen, S., Choat, B., Hacke, U.G., Pratt, R.B., Bhaskar, R., Brodribb, T.J., Bucci, S.J., Cao, K.-F., Cochard, H., Delzon, S., Domec, J.-C., Fan, Z.-X., Feild, T.S., Jacobsen, A.L., Johnson, D.M., Lens, F., Maherali, H., Martínez-Vilalta, J., Mayr, S., McCulloh, K.A., Mencuccini, M., Mitchell, P.J., Morris, H., Nardini, A., Pittermann, J., Plavcová, L., Schreiber, S.G., Sperry, J.S., Wright, I.J., Zanne, A.E., 2016. Weak tradeoff between xylem safety and xylem-specific hydraulic efficiency across the world’s woody plant species. New Phytol. 209, 123–136. https://doi.org/10.1111/nph.13646

24. Hacke, U.G., Jansen, S., 2009. Embolism resistance of three boreal conifer species varies with pit structure. New Phytol. 182, 675–686. https://doi.org/10.1111/j.1469-8137.2009.02783.x

25. Hacke, U.G., Sperry, J.S., Pittermann, J., 2004. Analysis of circular bordered pit function - II. Gymnosperm tracheids with torus-margo pit membranes. Am. J. Bot. 91, 386–400.

26. Hammond, W.M., 2020. A matter of life and death: Alternative stable states in trees, from xylem to ecosystems. Front. For. Glob. Chang.

27. Johnson, D.M., Katul, G., Domec, J.-C., 2022. Catastrophic hydraulic failure and tipping points in plants. Plant. Cell Environ. 45, 2231–2266. https://doi.org/10.1111/pce.14327

28. Jyske, T., Hölttä, T., 2015. Comparison of phloem and xylem hydraulic architecture in Picea abies stems. New Phytol. 205, 102–115. https://doi.org/10.1111/nph.12973

29. Kiorapostolou, N., Petit, G., 2019. Similarities and differences in the balances between leaf, xylem and phloem structures in *Fraxinus ornus* along an environmental gradient 39, 234–242. https://doi.org/10.1093/treephys/tpy095

30. Koçillari, L., Olson, M.E., Suweis, S., Rocha, R.P., Lovison, A., Cardin, F., Dawson, T.E., Echeverría, A., Fajardo, A., Lechthaler, S., Martínez-Pérez, C., Marcati, C.R., Chung, K.-F., Rosell, J.A., Segovia-Rivas, A., Williams, C.B., Petrone-Mendoza, E., Rinaldo, A., Anfodillo, T., Banavar, J.R., Maritan, A., 2021. The Widened Pipe Model of plant hydraulic evolution. Proc. Natl. Acad. Sci. 118, e2100314118. https://doi.org/10.1073/pnas.2100314118

31. Kolb, K.J., Sperry, J.S., 1999. Differences in drought adaptation between subspecies of sagebrush (*Artemisia tridentata*). Ecology 80, 2373–2384.

32. Lazzarin, M., Crivellaro, A., Williams, C.B., Dawson, T.E., Mozzi, G., Anfodillo, T., 2016. Tracheid and pit anatomy vary in tandem in a tall Sequoiadendron giganteum tree. IAWA J. 37, 172–185. https://doi.org/10.1163/22941932-20160129

33. Lechthaler, S., Kiorapostolou, N., Pitacco, A., Anfodillo, T., Petit, G., 2020. The total path length hydraulic resistance according to known anatomical patterns: What is the shape of the root-to-leaf tension gradient along the plant longitudinal axis? J. Theor. Biol. 502, 110369. https://doi.org/10.1016/j.jtbi.2020.110369

34. Lechthaler, S., Turnbull, T., Gelmini, Y., Pirotti, F., Anfodillo, T., Adams, M.A., Petit, G., 2019. A standardization method to disentangle environmental information from axial trends of xylem anatomical traits. Tree Physiol. 39, 495–502.

35. Lens, F., Gleason, S.M., Bortolami, G., Brodersen, C., Delzon, S., Jansen, S., 2022. Functional xylem characteristics associated with drought-induced embolism in angiosperms. New Phytol. 236, 2019–2036. https://doi.org/10.1111/nph.18447

36. Lintunen, A., Kalliokoski, T., 2010. The effect of tree architecture on conduit diameter and frequency from small distal roots to branch tips in Betula pendula, Picea abies and Pinus sylvestris. Tree Physiol. 30, 1433–1447. https://doi.org/10.1093/treephys/tpq085

37. Lintunen, A., Kalliokoski, T., Niinemets, Ü., 2010. The effect of tree architecture on conduit diameter and frequency from small distal roots to branch tips in *Betula pendula*, *Picea abies* and *Pinus sylvestris*. Tree Physiol. 30, 1433–1447. https://doi.org/10.1093/treephys/tpq085

38. McMahon, T.A., Kronauer, R.E., 1976. Tree structures - deducing principle of mechanical design. J. Theor. Biol. 59, 443–466.

39. Niklas, K.J., 1995. Size-dependent allometry of tree height, diameter and trunk taper. Ann. Bot. 75, 217–227.

40. Olson, M.E., Soriano, D., Rosell, J.A., Anfodillo, T., Donoghue, M.J., Edwards, E.J., León-Gómez, C., Dawson, T., Camarero Martínez, J.J., Castorena, M., Echeverría, A., Espinosa, C.I., Fajardo, A., Gazol, A., Isnard, S., Lima, R.S., Marcati, C.R., Méndez-Alonzo, R., 2018. Plant height and hydraulic vulnerability to drought and cold. Proc. Natl. Acad. Sci. 115, 7551–7556. https://doi.org/10.1073/pnas.1721728115

41. Petit, G., Anfodillo, T., 2009. Plant physiology in theory and practice: An analysis of the WBE model for vascular plants. J. Theor. Biol. 259, 1–4. https://doi.org/10.1016/j.jtbi.2009.03.007

42. Petit, G., Anfodillo, T., Carraro, V., Grani, F., Carrer, M., 2011. Hydraulic constraints limit height growth in trees at high altitude. New Phytol. 189, 241–252. https://doi.org/10.1111/j.1469-8137.2010.03455.x

43. Petit, G., Anfodillo, T., Mencuccini, M., 2008. Tapering of xylem conduits and hydraulic limitations in sycamore (*Acer pseudoplatanus*) trees. New Phytol. 177, 653–664. https://doi.org/10.1111/j.1469-8137.2007.02291.x

44. Petit, G., Mencuccini, M., Carrer, M., Prendin, A.L., Höltta, T., 2023. Axial conduit widening, tree height and height growth rate set the hydraulic transition of sapwood into heartwood. J. Exp. Bot.

45. Petit, G., Pfautsch, S., Anfodillo, T., Adams, M.A., 2010. The challenge of tree height in *Eucalyptus regnans*: When xylem tapering overcomes hydraulic resistance. New Phytol. 187, 1146–1153. https://doi.org/10.1111/j.1469-8137.2010.03304.x

46. Pfautsch, S., Aspinwall, M.J., Drake, J.E., Chacon-Doria, L., Langelaan, R.J.A., Tissue, D.T., Tjoelker, M.G., Lens, F., 2018. Traits and trade-offs in whole-tree hydraulic architecture along the vertical axis of Eucalyptus grandis. Ann. Bot. 121, 129–141. https://doi.org/10.1093/aob/mcx137

47. Pittermann, J., Sperry, J.S., Hacke, U.G., Wheeler, J.K., Sikkema, E.H., 2006. Inter-tracheid pitting and the hydraulic efficiency of conifer wood: The role of tracheid allometry and cavitation protection. Am. J. Bot. 93, 1265–1273.

48. Prendin, A.L., Petit, G., Carrer, M., Fonti, P., Björklund, J., Von Arx, G., 2017. New research perspectives from a novel approach to quantify tracheid wall thickness. Tree Physiol. 37, 1–8. https://doi.org/10.1093/treephys/tpx037

49. Prendin, A.L., Petit, G., Fonti, P., Rixen, C., Dawes, M.A., von Arx, G., 2018. Axial xylem architecture of *Larix decidua* exposed to CO2enrichment and soil warming at the tree line. Funct. Ecol. 32, 273–287. https://doi.org/10.1111/1365-2435.12986

50. R Development Core Team, 2022. A language and environment for statistical computing. R Foundation for Statistical Computing, Wien.

51. Rosner, S., Gierlinger, N., Klepsch, M., Karlsson, B., Evans, R., Lundqvist, S.O., Světlík, J., Børja, I., Dalsgaard, L., Andreassen, K., Solberg, S., Jansen, S., 2018. Hydraulic and mechanical dysfunction of Norway spruce sapwood due to extreme summer drought in Scandinavia. For. Ecol. Manage. 409, 527–540. https://doi.org/10.1016/j.foreco.2017.11.051

52. Rosner, S., Heinze, B., Savi, T., Dalla-Salda, G., 2019a. Prediction of hydraulic conductivity loss from relative water loss: new insights into water storage of tree stems and branches. Physiol. Plant. 165, 843–854. https://doi.org/10.1111/ppl.12790

53. Rosner, S., Johnson, D.M., Voggeneder, K., Domec, J.-C., 2019b. The conifer-curve: fast prediction of hydraulic conductivity loss and vulnerability to cavitation. Ann. For. Sci. 76, 82. https://doi.org/10.1007/s13595-019-0868-1

54. Rosner, S., Nöbauer, S., Voggeneder, K., 2021. Ready for screening: Fast assessable hydraulic and anatomical proxies for vulnerability to cavitation of young conifer sapwood. Forests. https://doi.org/10.3390/f12081104

55. Savi, T., Casolo, V., Dal Borgo, A., Rosner, S., Torboli, V., Stenni, B., Bertoncin, P., Martellos, S., Pallavicini, A., Nardini, A., 2019. Drought-induced dieback of Pinus nigra: a tale of hydraulic failure and carbon starvation. Conserv. Physiol. 7, coz012. https://doi.org/10.1093/conphys/coz012

56. Schenk, H.J., Espino, S., Visser, A., Esser, B.K., 2016. Dissolved atmospheric gas in xylem sap measured with membrane inlet mass spectrometry. Plant. Cell Environ. 39, 944–950. https://doi.org/10.1111/pce.12678

57. Schulte, P.J., 2012. Vertical and radial profiles in tracheid characteristics along the trunk of Douglas-fir trees with implications for water transport. Trees 26, 421–433. https://doi.org/10.1007/s00468-011-0603-5

58. Song, Y., Poorter, L., Horsting, A., Delzon, S., Sterck, F., 2022. Pit and tracheid anatomy explain hydraulic safety but not hydraulic efficiency of 28 conifer species. J. Exp. Bot. 73, 1033–1048. https://doi.org/10.1093/jxb/erab449

59. Sperry, J.S., Hacke, U.G., Pittermann, J., 2006. Size and function in conifer tracheids and angiosperm vessels. Am. J. Bot. 93, 1490–1500.

60. Spicer, R., Gartner, B.L., 2001. The effects of cambial age and position within the stem on specific conductivity in Douglas-fir (Pseudotsuga menziesii) sapwood. Trees 15, 222–229. https://doi.org/10.1007/s004680100093

61. Tyree, M.T., Zimmermann, M.H., 2002. Xylem structure and the ascent of sap, Springer Series in Wood Science. Springer, Berlin, Heidelberg. https://doi.org/10.1007/978-3-662-04931-0

62. von Arx, G., Carrer, M., 2014. ROXAS - a new tool to build centuries-long tracheid-lumen chronologies in conifers. Dendrochronologia 32, 290–293. https://doi.org/10.1016/j.dendro.2013.12.001

63. West, G.B., Brown, J.H., Enquist, B.J., 1999. A general model for the structure and allometry of plant vascular systems. Nature 400, 664–667. https://doi.org/10.1038/23251

64. West, G.B., Brown, J.H., Enquist, B.J., 1997. A general model for the origin of allometric scaling laws in biology. Science (80-.). 276, 122–126.

65. Zar, J., 1999. Biostatistical analysis, New Jersey. Prentice Hall, Upper Saddle River.

66. Zuur, A.F., Ieno, E.N., Walker, N.J., Saveliev, A.A., Smith, G.M., 2009. Reviewer: Aaron Christ Alaska Department of Fish and Game Mixed Effects Models and Extensions in Ecology with R. JSS J. Stat. Softw. 32, 2–4.

